# A New Labelling Method of ^99m^Tc-PSMA-HBED-CC

**DOI:** 10.1101/2024.10.31.621442

**Authors:** Benchamat Phromphao, Shuichi Shiratori

## Abstract

**Objective:** ^68^Ga-PSMA-HBED-CC (^68^Ga-PSMA-11) was approved by the US FDA as the first PSMA-targeted PET imaging drug for patients with prostate cancer. However, the utility of ^68^Ga-PSMA-HBED-CC may be limited due to PET/CT or PET/MR accessibility and ^68^GaCl_3_ availability produced from ^68^Ge/^68^Ga generator or cyclotron. Thus, in-house preparation of ^99m^Tc-PSMA-HBED-CC was developed as an alternative to ^68^Ga-PSMA-HBED-CC to be ubiquitous and affordable in the worldwide population.

**Methods:** A solution of ^99m^Tc-pertechnetate was added to PSMA-HBED-CC and 4% SnCl_2_·2H_2_O in a 10 mL sterile vial. The mixture was heated at 100°C for 15 minutes and then allowed to cool to room temperature. Labeling conditions were optimized to maximize the radiochemical yield of ^99m^Tc-PSMA-HBED-CC. The chelation completeness was monitored using instant thin layer chromatography (iTLC), and the stability of ^99m^Tc-PSMA-HBED-CC was subsequently evaluated.

**Results:** The radiolabeling of ^99m^Tc-PSMA-HBED-CC was succeeded using the appropriate amount of 10 µg PSMA-HBED-CC 3 µg SnCl_2_·2H_2_O and ^99m^Tc-pertechnetate 370 MBq at 100 °C for 15 mins, yielded the best result in high radiochemical yield (71.49 ± 2.42%), radiochemical purity (98.29 ± 2.65%), specific activity of 37.84 ± 1.47 GBq/µmol. ^99m^Tc-PSMA-HBED-CC is stable with radiochemical purity of more than 95% within 4 hrs at room temperature.

**Conclusion:** A new labelling method of ^99m^Tc-PSMA-HBED-CC was developed. Quality control parameters of ^99m^Tc-PSMA-HBED-CC met the criteria in accordance with the European Pharmacopoeia.

## Introduction

Prostate cancer is the most common malignancy found in men and the second leading cause of cancer death worldwide.^(1)^ Over the past two decades, the initial diagnosis and follow-up have been serum prostate-specific antigen (PSA) levels, digital rectal examination, and some conventional imaging techniques including ultrasound, CT, MRI, and bone scintigraphy but none provides highly specific and sensitive detection. Although transrectal ultrasound-guided (TRUS) prostate biopsy is currently accepted as the gold standard to provide the histopathological diagnosis of prostate cancer,^(2)^ it is an invasive procedure that resulted in a risk of side effects and the accuracy of diagnosis. The accurate definition of tumor burden and its staging is particularly important for effective treatment selection. Therefore, molecular imaging, a non-invasive method, has been employed to visualize the tumor in both soft tissue and bone with higher specific and sensitive detection, monitored response to therapy to improve management of prostate cancer, clinical outcome, and patient’s quality of life.

The Food and Drug Administration (FDA) approved ^68^Ga-PSMA-HBED-CC (^68^Ga-PSMA-11, previous name: ^68^Ga-DKFZ-PSMA-11, generic name: ^68^Ga-Gazetotide) as the first PSMA-targeted positron emission tomography (PET) imaging drug for men with prostate cancer.^(3)^ The development of ^68^Ga-PSMA-HBED-CC which targets prostate-specific membrane antigen (PSMA) has offered new perspectives for prostate cancer diagnosis and evaluation of therapeutic response.^(4)^ PSMA is a cell surface transmembrane protein with 750 amino acids type II glycoprotein that primarily expresses in normal prostate epithelium and is overexpressed in prostate cancer cells including bone metastasis.^(5)^ X-ray crystal structure analysis of PSMA, also known as *N*-acetylated *L*-aspartyl-*L*-glutamate peptidase (NAALADase I), has identified the critical interaction of potent inhibitors within the hydrophobic active site of the enzyme.^(6)^ Consequently, several classes of NAALADase I inhibitors had been exploited for structure-based design platforms, leading to the novel synthesis of PSMA-HBED-CC.^(7-9)^ To date, ^68^Ga-PSMA-HBED-CC has explicitly demonstrated superior detection of PSMA-positive prostate cancer lesions in recurrent and metastatic sites over conventional imaging methods^(10, 11)^ and two other FDA-approved PET tracers, ^18^F-Fluciclovine, ^11^C-Choline that used in patients with suspected cancer recurrence.^(12)^ Recently, FDA also approved ^18^F-Piflufolastat as the second PSMA-targeted positron emission tomography (PET) imaging drug with prostate cancer.^(13)^

Besides ^68^Ga-PSMA-HBED-CC, the preferential use of ^99m^Tc-labeled urea-based PSMA inhibitor has received interest as an alternative option to widespread the advantages of PSMA imaging due to a number of prostate cancer patients who are scheduled on PSMA imaging. The hybrid modality of SPECT/CT offers a wide range of workhorses in nuclear medicine with lower financial access, especially the remote medical center in which PET/CT facility is not available. Although the spatial resolution of ^99m^Tc is not as good as that of ^68^Ga, ^99m^Tc provides a sufficiently long half-life of 6 hours in both preparation and accumulation in the target site. Moreover, the decay range of ^99m^Tc is short enough to minimize radiation exposure to patients and medical staff.

While some ^99m^Tc-labeled PSMA tracers have been previously reported using various PSMA ligands with several forms of complexation for example [^99m^Tc(CO)_3_(L)_3_]^+, (14-16)^ MAG3 based ^99m^Tc-PSMA-I&S,^(17) 99m^Tc-MIP,^(18-21) 99m^Tc-HYNIC-PSMA,^(22, 23)^ peptide-chelator based ^99m^Tc-DUPA,^(24) 99m^Tc-PSMA-T4, ^(25) 99m^Tc-PSMA-tricarbonyl-HBED-CC^(26)^ and ^99m^Tc-PSMA-HBED-CC,^(27)^ it challenges to develop a convenient labelling method in a single step without co-ligand for ^99m^Tc-complexation. According to our experience in theranostics, we adapted the routine standard labeling procedure of ^68^Ga-PSMA-HBED-CC with the rationale that HBED-CC would serve as a suitable chelator in a mimic manner to DTPA as shown in Fig 1. Our attention focused on optimizing the labeling parameters to improve the radiosynthesis of ^99m^Tc-PSMA-HBED-CC.

**Fig 1.**
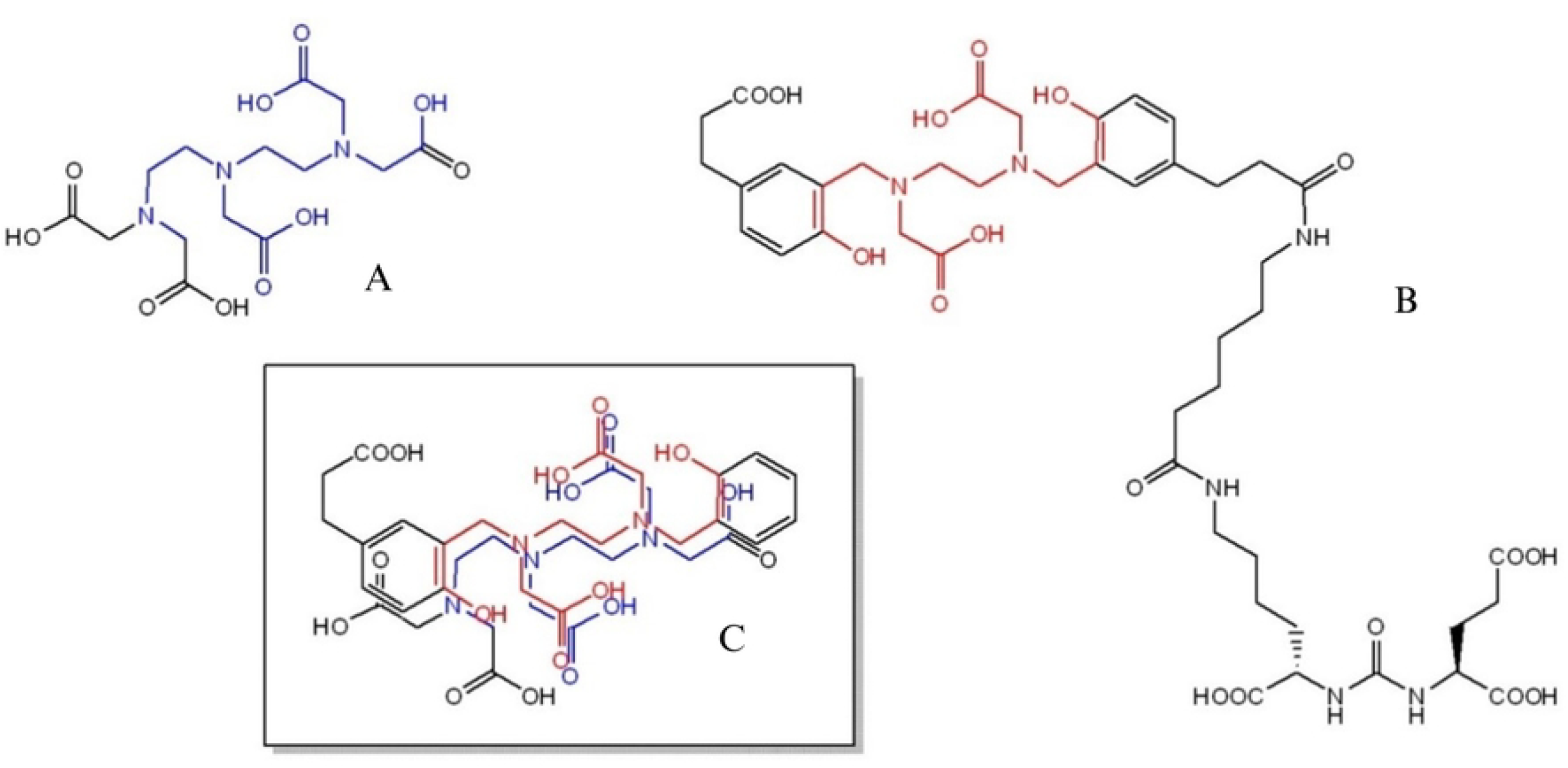
Chemical structures of (A) DTPA, (B) PSMA-HBED-CC and (C) imposed chelating motif.

## Materials and Methods

PSMA-HBED-CC was purchased from ABX advanced biochemical compounds (GmbH, Germany). Sodium ^99m^Tc-pertechnetate was purchased from Global Medical Solution (Thailand). Stannous chloride dihydrate and hydrochloric acid were purchased from Sigma-Aldrich (Germany). The stock solution of 4% stannous chloride was freshly prepared and kept in refrigerator. All chemicals and solvents were used without further purification unless otherwise noted. The C18 cartridge was purchased from Waters (USA). The TLC scanner Raytest model MiniGita was used.

### Preparation of 4% stannous chloride

SnCl_2_·2H_2_O 0.144 mg was added to concentrate HCl 0.75 mL, followed by heating at 100 °CC for 5 minutes. After cooling down to room temperature, 6 N HCl 2.25 mL was added to be 4% SnCl_2_·2H_2_O. This reducing agent solution should be freshly prepared before labeling.

### Labelling of PSMA-HBED-CC with Tc-99m

To a mixture of 4% SnCl_2_·2H_2_O solution calculated as amount of SnCl_2_ and an aliquot of PSMA-HBED-CC (10 µg in H_2_O 100 µL) in 10 mL sterile vial, ^99m^Tc-pertechnetate 370 MBq was added. The labeling was performed at 100 °CC for 15 minutes in a heating block, followed by a 10-minute cool down to reach room temperature. The crude product was passed through a C18 cartridge. ^99m^Tc-PSMA-HBED-CC was slowly purged from a C18 cartridge using EtOH:H_2_O (1:1) 2 mL to the final product vial.

### Quality control

Radiochemical purity (RCP) analyses were performed using instant thin-layer chromatography (iTLC) on silica paper strips as a stationary phase with two different mobile phases. The free form of ^99m^TcO_4_^-^ was determined using 0.9% normal saline, whereas ^99m^Tc-colloid formation was determined using acetone as the mobile phase. The radioactivity distribution on iTLC strips was also determined on a TLC scanner. Subsequently, radiochemical yield (RCY) was calculated. The pH of the final product was determined using a pH indicator.

### Stability test

The chemical stability of ^99m^Tc-PSMA-HBED-CC was carried out by incubating at room temperature for 6 hours and monitored by iTLC every hour.

## Results

### Labelling of 99mTc-PSMA-HBED-CC

The labeling parameters were investigated to achieve the highest possible radiochemical yield. The quantity of PSMA-HBED-CC used in each experiment remained constant at 10 µg (0.011 µmol). In accordance with the DTPA cold kit formulation, the appropriate radioactivity of ^99m^Tc-pertechnetate was determined in direct correlation with approximate 370 MBq (10 mCi), while maintaining the solution’s pH at 5.0, as indicated in Table 1.

**Table 1.**
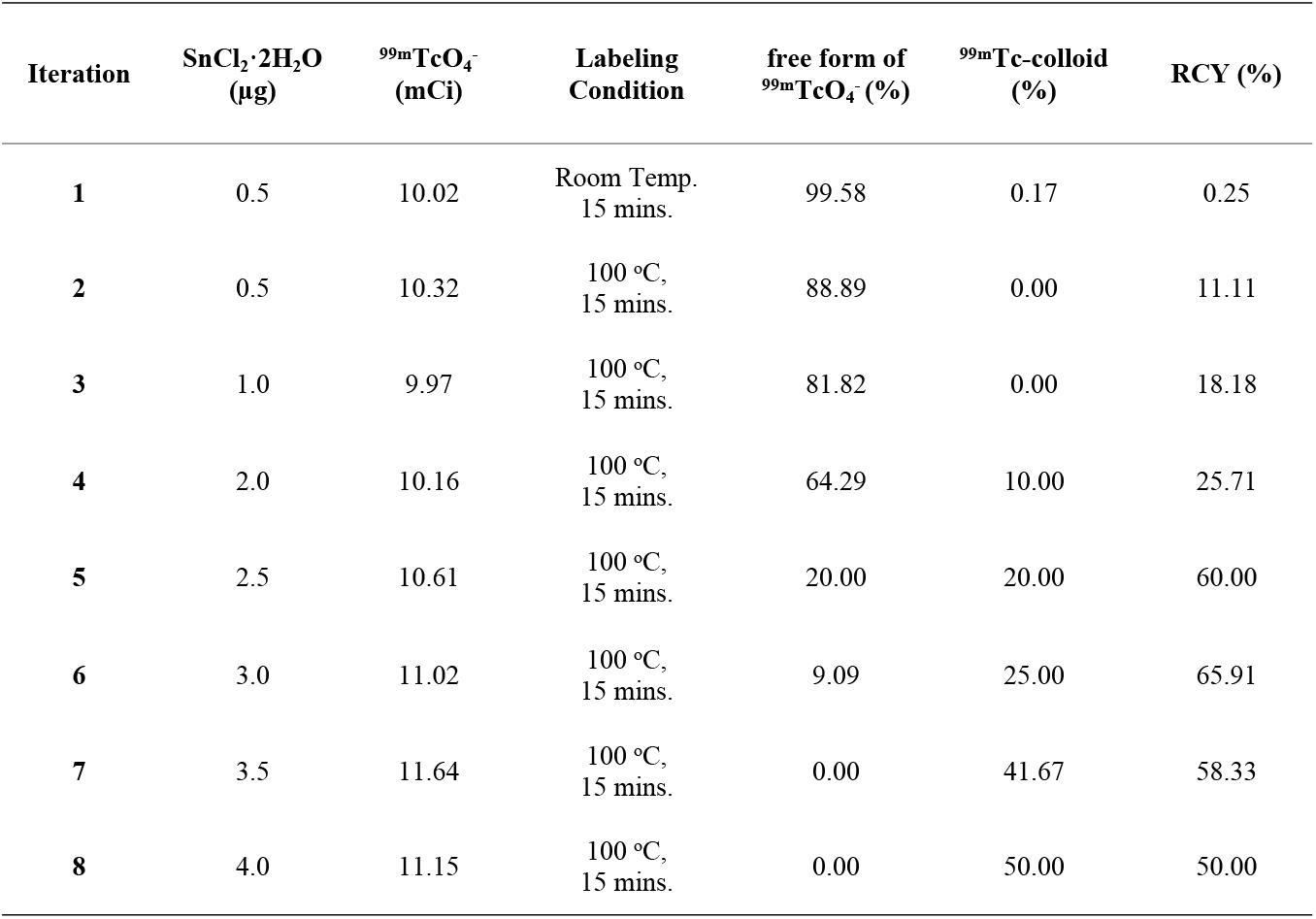
Labeling parameters in 99mTc-PSMA-HBED-CC

In the preliminary experiment, 10.02 mCi of ^99m^TcO_4_^-^ was combined with 0.5 µg of SnCl_2_ at room temperature for 15 minutes resulted in a RCY of 0.25%, with 99.58% of ^99m^TcO_4_^-^ remaining unbound. Subsequently, the reaction temperature was increased to 100°C for 15 minutes, in alignment with the standard procedure for labeling ^68^Ga-PSMA-HBED-CC. This adjustment resulted in a RCY of 11.11%, affirming the choice to set the reaction conditions at 100°C for 15 minutes. In iterations 3 to 8, the amount of SnCl_2_ was progressively adjusted to enhance the RCY. By increasing the SnCl_2_ quantity by 0.5 µg in each iteration, the highest RCY of 65.91% was achieved in the sixth iteration. However, when the amount of SnCl_2_ exceeded 3.5 µg, complete chelation occurred between PSMA-HBED-CC and ^99m^TcO_4s_^-^, which also led to a significant rise in the formation of hydrolyzed species, ultimately diminishing the RCY.

### Chemical stability of 99mTc-PSMA-HBED-CC

The chemical stability of ^99m^Tc-PSMA-HBED-CC was evaluated using the optimized labelling method in 6^th^ iteration that produced the highest RCY. The compound was incubated at room temperature for 6 hours, and RCP was checked hourly through iTLC (Fig 2). The findings are shown in Fig 3. In general, the RCP of Tc-99m radiopharmaceuticals should meet or exceed 95%. This study demonstrated that the RCP of ^99m^Tc-PSMA-HBED-CC remained above 95% for up to 4 hours after labelling, and stayed above 90% at the 5th and 6th hours.

**Fig 2.**
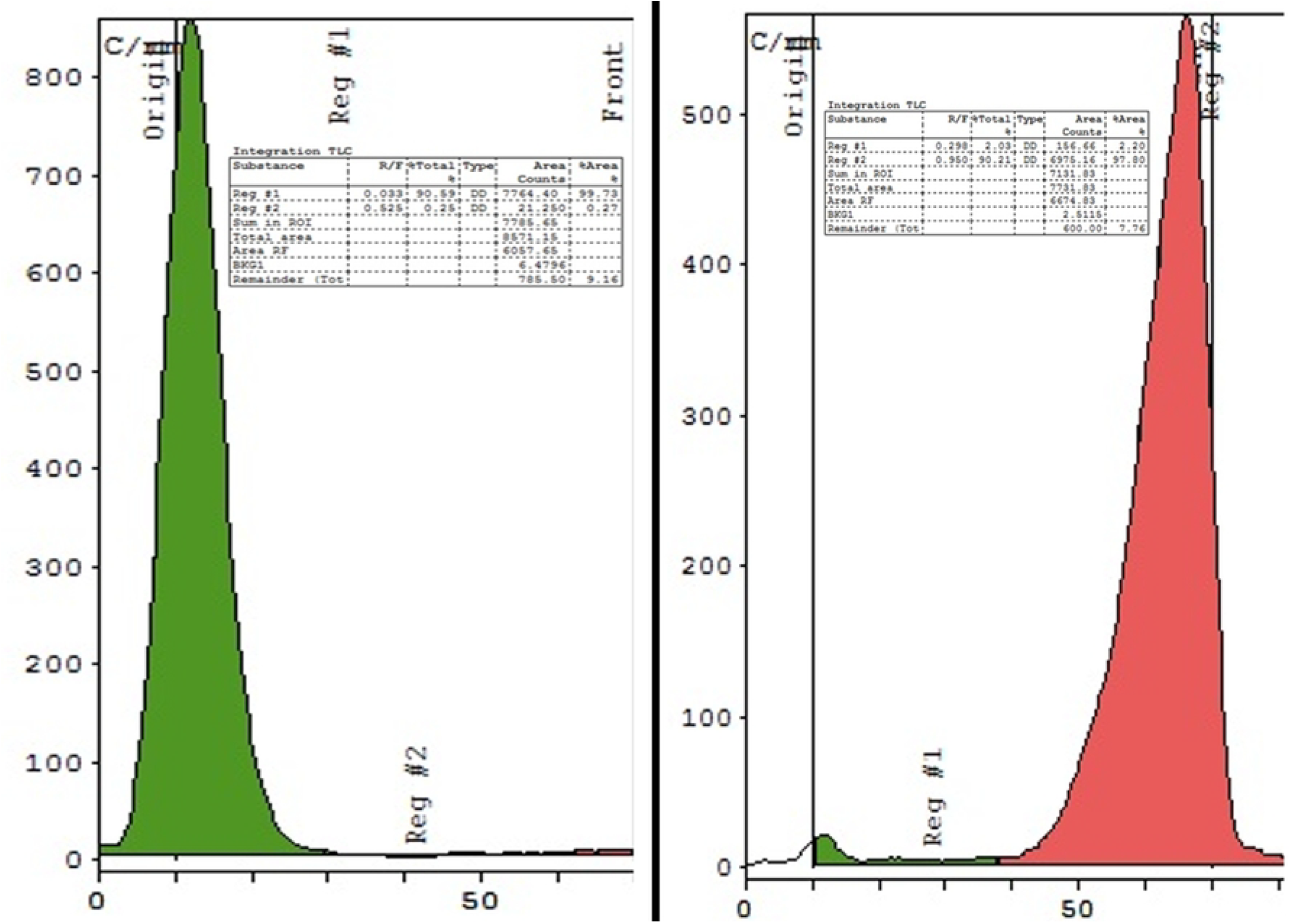
iTLC Chromatogram of ^99m^Tc-PSMA-HBED-CC; (left) in acetone as mobile phase, (right) in 0.9% normal saline as mobile phase.

**Fig 3.**
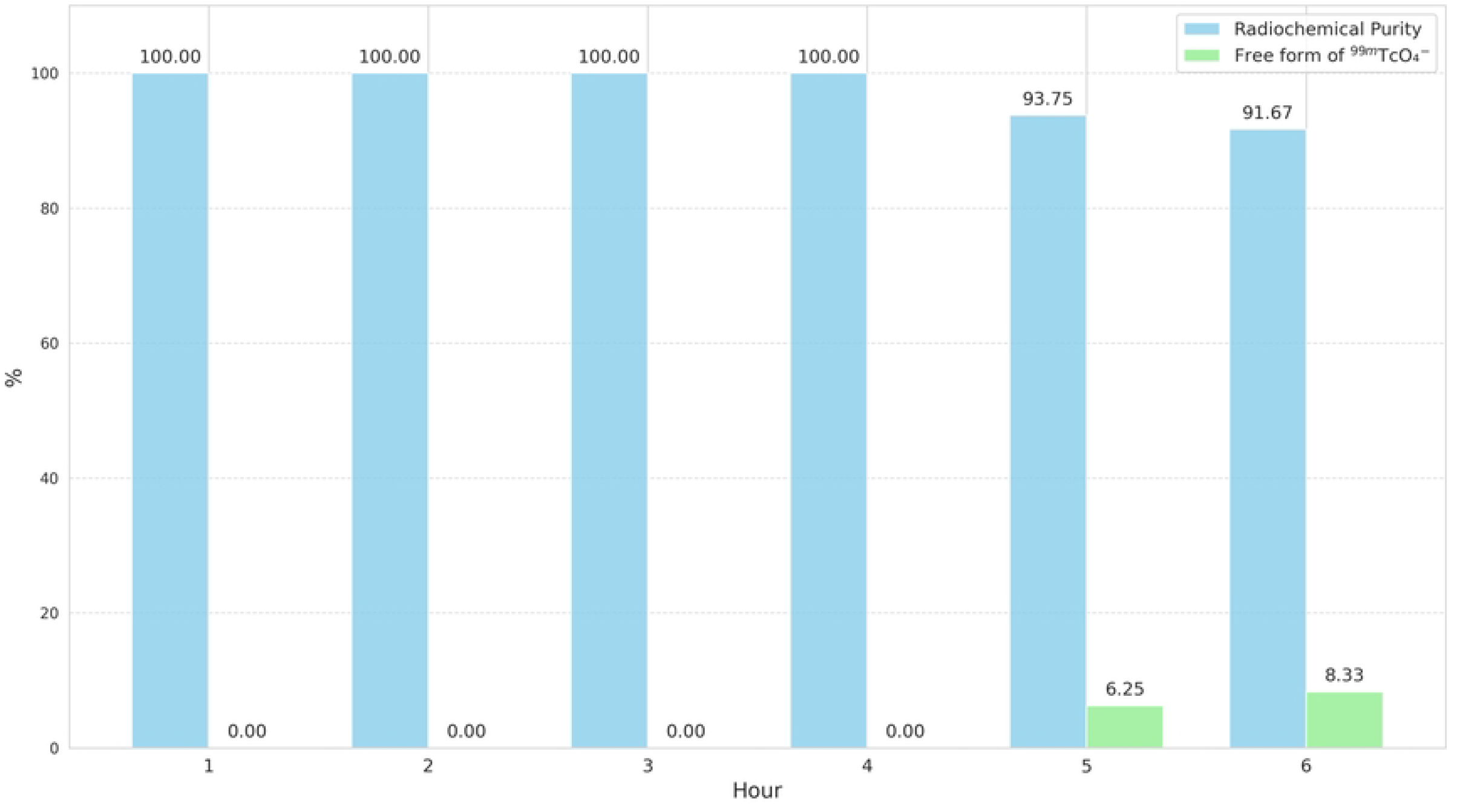
Chemical stability of ^99m^Tc-PSMA-HBED-CC at room temperature

## Discussion

Since the identification of PSMA as an antigen and the discovery of the specific antibody 7E11-C5 (Capromab) for both normal and malignant prostate epithelium, as reported by Murphy and Horoszewicz et al.,^(28)^ PSMA has emerged as a crucial target for prostate cancer cells.^(29-31)^ In 2012, Eder et al. developed the urea-based PET tracer ^68^Ga-PSMA-HBED-CC (formerly ^68^Ga-DKFZ-PSMA-11) to address the limitations of the lead antibody J591.^(32)^ The U.S. Food and Drug Administration approved ^68^Ga-PSMA-HBED-CC as the first PET imaging agent for PSMA-positive lesions in men with prostate cancer in December 2020^(3)^ and ^18^F-Piflufolastat as the second PET imaging agent for PSMA-positive lesions in men with prostate cancer in May 2021^(13)^ Inspired by this breakthrough, our study aimed to develop ^99m^Tc-PSMA-HBED-CC as a SPECT imaging analogue to increase accessibility for prostate cancer diagnosis.

PSMA-HBED-CC was chosen for this study due to its widespread clinical use and commercial availability. The dose of PSMA-HBED-CC was standardized at 10 μg. To optimize the manual labelling of ^99m^Tc-PSMA-HBED-CC, various labelling conditions and amounts of calculated SnCl_2_ were assessed. The results, presented in Table 1, indicate that ^99m^Tc-PSMA-HBED-CC is a thermodynamically favorable product. Using 2.5-4.0 μg of SnCl_2_ resulted in a RCY exceeding 50%. When the SnCl_2_ amount exceeded 3.5 μg, no uncomplexed Tc-99m was detected, indicating effective reduction of TcO_4_^-^. However, higher amounts of SnCl_2_ also increased colloid formation. Optimal quantitative radiolabelling of 10 μg of PSMA-HBED-CC was achieved with 3.0 μg of SnCl_2_. To prevent colloid formation, both SnCl_2_ and PSMA-HBED-CC must be present in the reaction mixture before adding TcO_4_^-^ to form the desired complex.

The chemical stability of ^99m^Tc-PSMA-HBED-CC was evaluated by incubating it at room temperature. As shown in Fig 2, it retained RCP above 95% for up to 4 hours. After this period, free Tc-99m increased. Therefore, it is recommended to use ^99m^Tc-PSMA-HBED-CC within 4 hours of preparation or store it in a refrigerator to maintain stability.

Vats et al.^(26)^ previously reported the preparation of ^99m^Tc-PSMA-HBED-CC using 50 mg of PSMA-HBED-CC, 40 mg of SnCl_2_, TcO_4_^-^ 740 MBq at pH 5, yielding a RCY of 60±5%, a RCP greater than 98%, and specific activity of 15±5 GBq/µmol. However, they did not conduct stability tests. Economically, our study used 10 µg of PSMA-HBED-CC, 3 µg of SnCl_2_, and 370 MBq of ^99m^Tc-pertechnetate at 100°C for 15 minutes, achieving a higher RCY (71.49±2.42%), RCP (98.29±2.65%), and specific activity (37.84±1.47 GBq/µmol). This method is more cost-effective and easier to manipulate.

## Conclusion

To optimize the labeling of PSMA-HBED-CC with ^99m^Tc-pertechnetate for prostate cancer imaging, the labeling procedure should be carried out at 100°C for 15 minutes using 3 µg of SnCl_2_, minimizing the presence of free ^99m^Tc-pertechnetate and colloid formation. Purification with a C18 cartridge is required to achieve RCP that complies with European Pharmacopoeia standards. The stability of ^99m^Tc-PSMA-HBED-CC remains robust for up to 4 hours at room temperature, with radiochemical purity exceeding 95%. It is recommended to use the labeled product within 4 hours of preparation.

## Declarations

Ethics approval and consent to participate: Not applicable.

## Consent for publication

Not applicable.

## Availability of data and material

Not applicable.

## Authors’ contributions

BP and SS contributed to the conceptualization, methodology, formulation, formal analysis, visualization. BP performed the kit manufacture. SS managed funding and acquisition. SS was a major contributor in writing the manuscript. All authors have read and agreed to the final version.

## Acknowledgements

Not applicable.

## Availability of data and material

Not applicable

